# Comparative genomics of two sequential *Candida glabrata* clinical isolates

**DOI:** 10.1101/132464

**Authors:** Luis Vale-Silva, Emmanuel Beaudoing, Van Du T. Tran, Dominique Sanglard

**Affiliations:** Institute of Microbiology, University of Lausanne and University Hospital Center, CH-1011 Lausanne (Switzerland); Center for Integrative Genomics, Lausanne Genomic Technologies Facility, CH-1015 Lausanne; Vital-IT Group, SIB Swiss Institute of Bioinformatics, CH-1015 Lausanne

**Keywords:** fungal pathogens, genome comparisons, drug resistance, adhesins

## Abstract

*Candida glabrata* is an important fungal pathogen which develops rapidly antifungal resistance in treated patients. It is known that azole treatments lead to antifungal resistance in this fungal species and that multidrug efflux transporters are involved in this process. Specific mutations in the transcriptional regulator *PDR1* result in upregulation of the transporters. In addition, we showed that the *PDR1* mutations can contribute to enhance virulence in animal models. We were interested in this study to compare genomes of two specific *C. glabrata* related isolates, one of which was azole-susceptible (DSY562) while the other was azole-resistant (DSY565). DSY565 contained a *PDR1* mutation (L280F) and was isolated after a time lapse of 50 days of azole therapy. We expected that genome comparisons between both isolates could reveal additional mutations reflecting host adaptation or even additional resistance mechanisms. The PacbBio technology used here yielded 14 major contigs (sizes 0.18 Mb-1.6 Mb) and mitochondrial genomes from both DSY562 and DSY565 isolates that were highly similar to each other. Comparisons of the clinical genomes with the published CBS138 genome indicated important genome rearrangements, but not between the clinical strains. Among unique features, several retrotransposons were identified in the genomes of the investigated clinical isolates. DSY562 and DSY565 contained each a large set of adhesin-like genes (101 and 107, respectively), which exceed by far the number of reported adhesins (66) in the CBS138 genome. Comparison between DSY562 and DSY565 yielded 17 non-synonymous SNPs (among which the expected *PDR1* mutation) as well as small size indels in coding regions (11) but mainly in adhesin-like genes. The genomes were containing a DNA mismatch repair allele of *MSH2* known to be involved in the so-called hypermutator phenotype of this yeast species and the number of accumulated mutations between both clinical isolates is consistent with the presence of a *MSH2* defect. In conclusion, this study is the first to compare genomes of *C. glabrata* sequential clinical isolates using the PacBio technology as an approach. The genomes of these isolates taken in the same patient at two different time points were exhibiting limited variations, even if submitted to the host pressure.

## Introduction

Infectious diseases caused by fungal pathogens represent a major threat to human health. It is estimated that about 2 billion people suffer several forms of fungal diseases worldwide, from which about 1.5 million deaths will occur every year (Brown *et al.* 2012). Invasive fungal infections cause on average 50% mortality, with total number of deaths comparable to tuberculosis and malaria (Brown *et al.* 2012). The most problematic fungal pathogens include *Candida*, *Aspergillus* and *Cryptococcus* species (Richardson and Lass-Flörl 2008). Within *Candida* species, *C. albicans* and *C. glabrata* are known as the most recovered species from infected patients (Marchetti *et al.* 2004). The current reports on the epidemiology of fungal diseases and their associated impact on human health reflect that the available therapeutic instruments, which include antifungal drugs, have limited efficacy. Only four major classes of antifungal drug are used to treat patients including azoles, polyenes, echinocandins and pyrimidine analogues. Each of these agents possess specific cellular targets and are declined in several formulations with different derivatives (Ruhnke 2014). In general, most of *Candida* species are susceptible to these drugs *in vitro*.

However, the use of antifungal agents has resulted in the inevitable occurrence of antifungal resistance. Antifungal resistance in a specific fungal isolate can be defined as a lack of activity of a given drug at concentrations that are higher than in a wild type isolate. Antifungal resistance has been reported virtually for all existing antifungal agents and major fungal pathogens (Perlin *et al.* 2015). Frequencies at which antifungal resistance occur in hospitalized patients are variable based on data collected by population-based surveillance programs and available for major fungal pathogens including *C. albicans* and *C. glabrata*. In general, antifungal resistance rates for *C. albicans* are low. In a study performed between 2008 and 2011, resistance to fluconazole (cut-off: ≥ 64 μg/ml) or echinocandins (cut-off: ≥4 μg/ml) ranged between 1-2% in bloodstream isolates (Cleveland *et al.* 2012). Resistance rates in *C. glabrata* are much higher: *C. glabrata* infections increased as a cause of invasive candidiasis from 18% of all BSI isolates in 1992-2001 to 25% in 2001-2007. Fluconazole resistance rates in *C. glabrata* increased from 9% to 14% at the same time (cut-off: ≥ 64 μg/ml) (Pfaller *et al.* 2003; 2009). Resistance of *C. glabrata* to the class of echinocandins is also increasing: within a 10-year period (2001–2010) in an US hospital, echinocandin resistance rate increased from 4.9 to 12.3% (Alexander *et al.* 2013). Similar trends are documented in Europe, however resistance rates are reported between 1-4% (Arendrup and Perlin 2014).

Recent studies have also documented the occurrence of multidrug resistance not only in *C. albicans* but also in *C. glabrata* (Sanglard 2015). *C. glabrata* cross-resistance between azoles and candins has captured more attention recently (Perlin 2015). For example, among echinocandin-resistant isolates sampled between 2008 and 2013 in two US surveillance hospital sites, 36% were also resistant to azoles (Pham *et al.* 2014). Between 2005 and 2013 in another US site, 10.3% *C. glabrata* isolates from cancer patients were resistant to caspofungin, from which about 60% were cross-resistant to azoles (Farmakiotis *et al.* 2014). Intriguingly, the later study reported that caspofungin exposure alone could induce multidrug resistance without azole exposure. While *C. glabrata* azole resistance is mediated principally by efflux-dependent mechanisms, candin resistance is the consequence of mutations in the genes (*FKS1*, *FKS2*) encoding the candin target enzyme (b-glucan synthase) (Vale-Silva and Sanglard 2015; Perlin 2015). It has been shown recently that *C. glabrata* can exhibit high mutation rates in the presence of antifungal agents. This property is based on the decreased activity of the DNA repair machinery that exists in specific clinical isolates (Healey, Zhao, *et al.* 2016). *C. glabrata* multidrug resistance is of concern when one considers that very few therapeutic alternatives are available.

We have shown in the past years that *C. glabrata* azole resistance is coupled with selective fitness advantages in animal models of infection (Ferrari *et al.* 2009; Vale-Silva *et al.* 2013). Specific isolates exhibit upregulation of the adhesin *EPA1* when the transcriptional activator responsible for azole resistance *CgPDR1* is mutated. When *EPA1* is upregulated, the consequence is that *C. glabrata* exhibits increased adherence to host cells, which is one of the factor that contributes to enhance virulence of this yeast species develops azole resistance (Vale-Silva *et al.* 2016). Our recent data also highlighted that *C. glabrata* can exhibit different degrees of adhesion to host cells. Given that adhesins are implicated in this process, it becomes relevant to address the repertoire of adhesins in specific *C. glabrata* lineages. Genome data are critical to achieve this goal, however the only available *C. glabrata* genome is up to now from the isolate CBS138 (Dujon *et al.* 2004). It is estimated that *C. glabrata* contains 63 ORF with adhesin properties (de Groot *et al.* 2013). In a recent study, we compared CBS138 to other clinical isolates and found that a specific isolate (DSY562) exhibited the much higher adherence capacities to epithelial cells as compared to CBS138. This result was not dependent on *EPA1* expression, since DSY562 expressed *EPA1* to much lower levels than CBS138.

These results stimulated our interest into the acquisition of the DSY562 genome. At the same time, it became also obvious to compare the genome of this isolate with one (DSY565) that acquired azole resistance within a time lapse of 50 days from DSY562 in a patient with oropharyngeal candidiasis (Sanglard *et al.* 1999). This way, evolutionary pressure on the derived isolate could be estimated at the level of genome differences.

Here we report genome sequences from both related isolates. We used several approach to resolve the genome with high accuracy. We analysed the occurrence of polymorphisms between the two strains and also addressed the repertoire of genes encoding adhesins.

## Results

### Genome data acquisition

*C. glabrata* DSY562 and DSY565 are paired clinical isolates that were isolated from a HIV-positive patient presenting oropharyngeal candidiasis. DSY565 was isolated after 50 days fluconazole therapy and exhibited azole resistance (Sanglard *et al.* 1999) (Table 1). The relationship between the two isolates was assessed at this time by RFLP and concluded a high relationship between the isolates. Before undertaking genome sequencing, both isolates were subjected to MLST typing according to Dodgson *et al.* (Dodgson *et al.* 2005) (Table 1) and thus confirmed the high relationship between both isolates.

**Table 1.** Characteristics of DSY562 and DSY565

| *C. glabrata*<br>clinical<br>strains | ST<sup>1)</sup> | MIC (µg/ml) |  |  | *PDR1*<br>mutation | Time<br>elapsed<br>(days) |
| --- | --- | --- | --- | --- | --- | --- |
|  |  | Fluconazole | Voriconazole | Amphotericin B |  |  |
| DSY562 | ST8 | 2 | 0.25 | 0.5 | - | - |
| DSY565 | ST8 | 64 | 1 | 0.5 | L280F | 50 |

<sup>1)</sup>: ST: sequence type

Genome sequencing was first approached by Illumina Hiseq 38 bp paired-end sequencing. Briefly, trimmed reads of both DSY5652 and DSY565 were aligned to the reference genome of CBS138 using the software CLC Genomics Workbench. In both cases, about 95% of reads aligned to the CBS genomes and coverage was 72-to 64-fold for DSY562 and DSY565 data, respectively. Details on comparisons can be found in supplementary **File S1** and **File S2.** Both comparisons yielded a high number of changes (85’711-85’986 including SNPs and indels) and resulted in about 10390 non-synonymous changes as compared to CBS138. Given that the genomes were aligned and compared directly to CBS138, they did not reflect actual genome rearrangements of strains DSY562 and DSY565. We therefore undertook an alternative genome sequencing approach using PacBio technologies enabling *de novo* assembly of large reads. Since this technology is prone to random intrinsic high error rates, we aimed to reach genome sequencing coverage between 200 and 300-fold, which therefore needed the use of several sequencing smrt cells. The PacBio data are summarized in Table 2. As summarized in this Table, the number of major assembled contigs (size > 20 kb) were 14 and 15 for DSY562 and DSY565, respectively. N50s were 1.06 and 1.17 Mb for both isolates, thus highlighting the high quality of the assemblies.

**Table 2:**
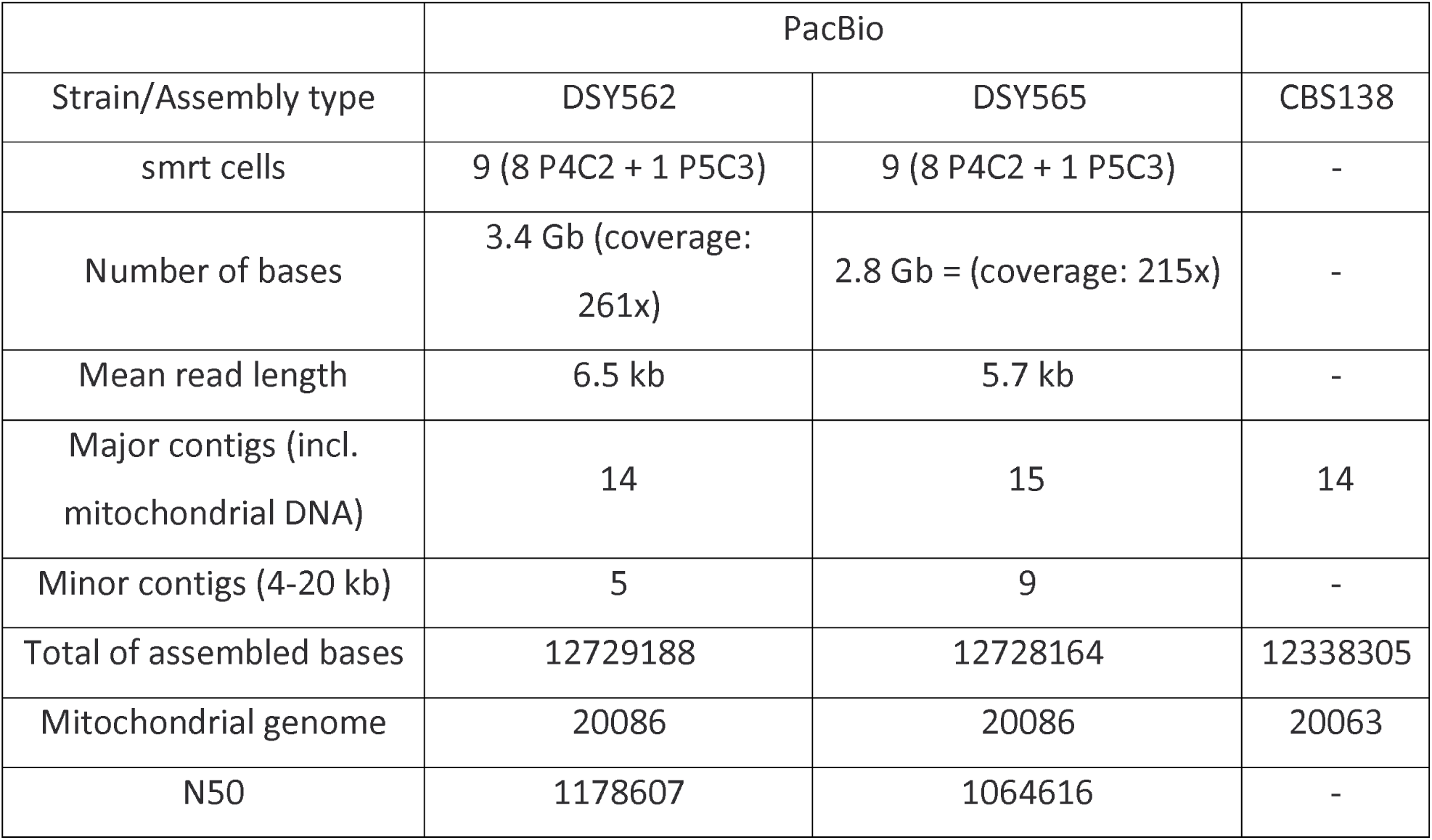
PacBio sequencing analysis of DSY562 and DSY565

### Genome comparisons between clinical strains and CBS138

Assembled contigs almost reconstituted the entire set of chromosomes that is known from the CBS138 genome. Fig. 1 shows the chromosome maps of DSY562 as compared to CBS138 as well as a circular map to highlight major translocations between the reference and the DSY562 genomes. For practical reasons, the naming of chromosomes (A to M) was kept as proposed initially for CBS138. The data highlight several chromosomal rearrangements between the two isolates. While Chr A, B, E, G, H and K kept similar structures between both strains, other chromosomes underwent several modifications. Chr C, D, J and M in DSY562 underwent exchanges from CBS138 chromosomes essentially at chromosome extremities. Parts of Chr L and I were added to Chr F (which was renamed as Chr F-L-I) in DSY562, thus yielding the largest chromosome in this species (1.59 Mb). Chr D and L were joined to the shortened Chr I (which was renamed as Chr I-L-D). The left arms of Chr L and Chr C were inverted as compared to CBS138. Remarkably is that the separate DSY565 assembly yielded the same chromosome arrangement as DSY562, with the exception that Chr J was not completely assembled in DSY565. This chromosome was still split in two parts (Chr J1 and J2, see Table 3) in DSY565. A CHEF was performed with strains CBS138 and DSY562/DSY565 to compare these assemblies with physical chromosome separations. Remarkably, size predictions from assemblies were corresponding well to physical separations (see Fig. 2). Red arrows shown in Fig. 2 correspond to sizes conservation of Chr A, B, G, H and J between the two strains, which is consistent with predictions. Chr C, D, E and F-L-I were shifted to higher sizes (yellow color) by CHEF in DSY562, which is also consistent with assembly predictions. Chr M and I-L-D exhibited decreased sizes (blue color) as compared to CBS138, which is also predicted from assemblies. Chr K from DSY562 showed discrepancy with expected patterns from CBS138, this result may however be due to intrinsic chromosome size variations observed by others for this type of strain (Bader *et al.* 2012).

**Figure 1.**
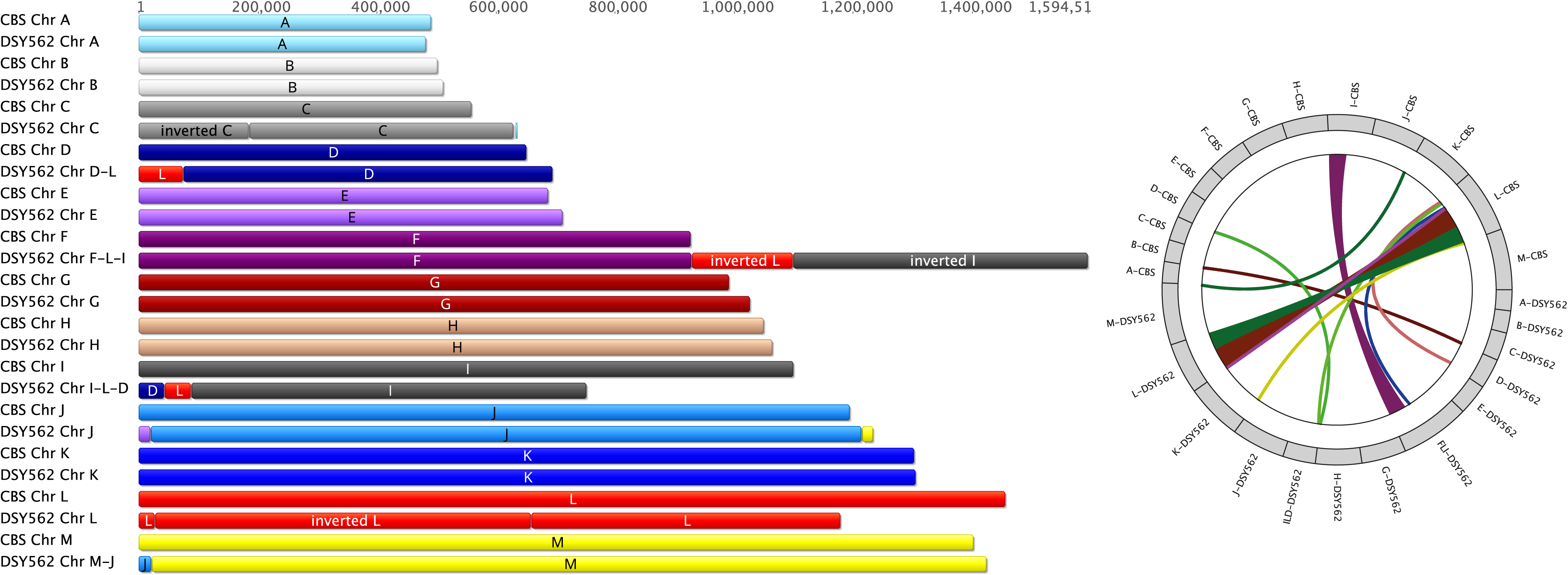
Chromosome structures of DSY562 as compared to CBS138. A circular map representing the major translocations between the CBS138 and DSY562 genomes was produced using J-Circos.v2 (An *et al.* 2015).

**Figure 2.**
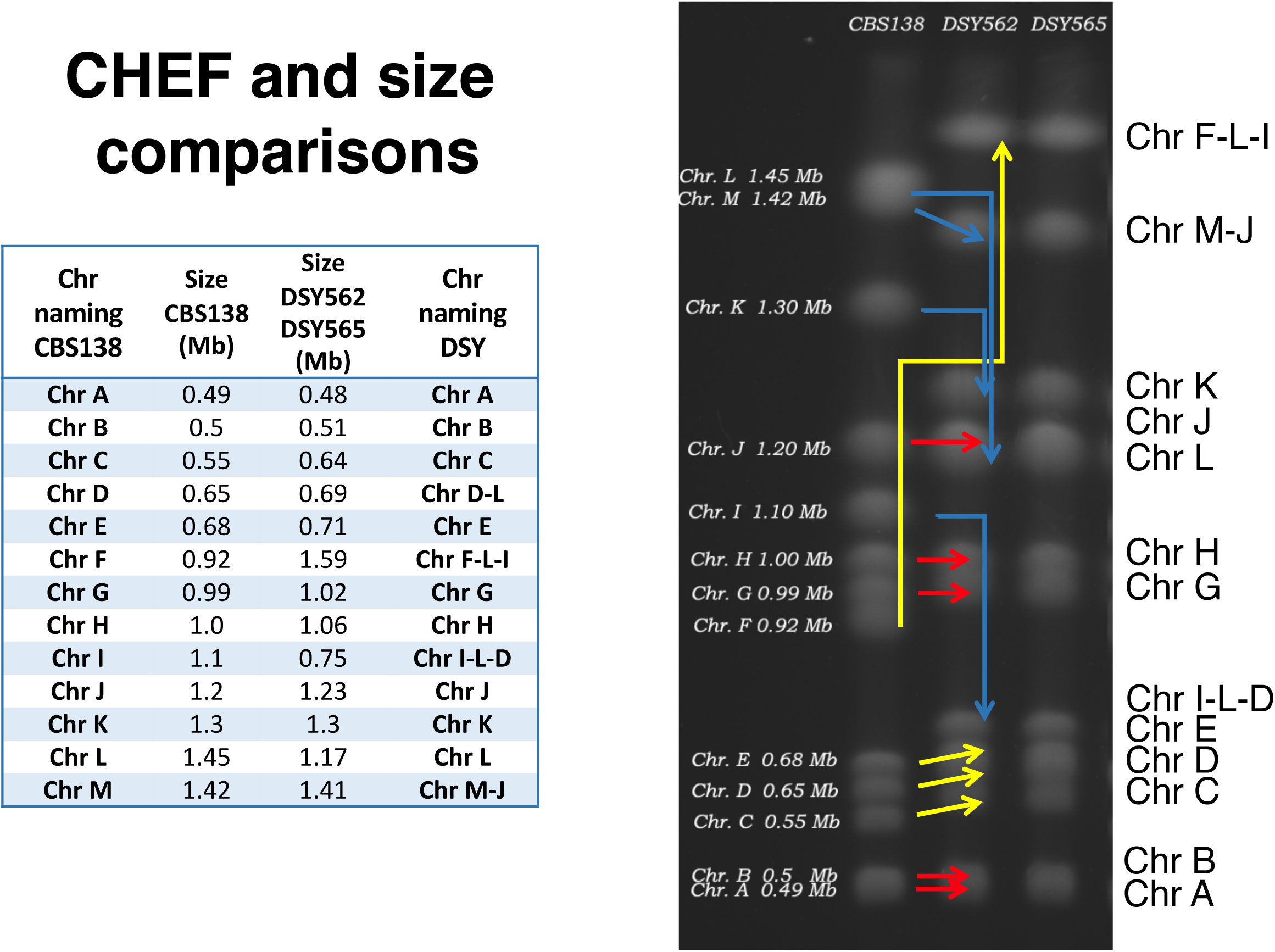
CHEF chromosome separations of isolates CBS138, DSY562 and DSY565. Chromosome sizes are indicated for CBS138. Colored arrows show the changes between the CBS and DSY562/DSY565 genomes. Red, yellow and blue colors signify size conservation, increased sizes and decreased sizes between the two different genomes. The attached table lists the chromosome sizes deduced from genome data.

**Table 3:** DSY562 and DSY565 annotation features

| DSY562 |  |  |  |  |  | DSY565 |  |  |  |  |  | CBS138 |  |  |  |  |
| --- | --- | --- | --- | --- | --- | --- | --- | --- | --- | --- | --- | --- | --- | --- | --- | --- |
| Chr | CDS | tRNA | rRNA | Introns | confirmed<br>Introns | Chr | CDS | tRNA | rRNA | Introns | confirmed<br>Introns | Chr | CDS | tRNA | rRNA | Introns |
| A | 201 | 9 |  | 10 | 7 | A | 201 | 9 |  | 10 | 6 | A | 205 | 9 |  | 9 |
| B | 218 | 9 |  | 4 | 3 | B | 218 | 9 |  | 4 | 3 | B | 220 | 9 |  | 4 |
| C | 242 | 7 |  | 4 | 1 | C | 240 | 7 |  | 4 | 1 | C | 235 | 7 |  | 7 |
| DL | 300 | 18 |  | 7 | 6 | DL | 300 | 18 |  | 7 | 7 | D | 288 | 18 |  | 6 |
| E | 279 | 9 |  | 8 | 5 | E | 279 | 9 |  | 8 | 5 | E | 282 | 9 |  | 10 |
| FLI | 661 | 29 |  | 19 | 15 | FLI | 661 | 8 |  | 19 | 15 | F | 387 | 15 |  | 12 |
| G | 438 | 18 |  | 18 | 16 | G | 438 | 29 |  | 18 | 17 | G | 440 | 18 |  | 18 |
| H | 470 | 14 |  | 11 | 9 | H | 470 | 18 |  | 11 | 9 | H | 466 | 14 |  | 12 |
| ILD | 278 | 18 |  | 2 | 2 | ILD | 278 | 14 |  | 2 | 2 | I | 470 | 26 |  | 6 |
| J | 522 | 17 |  | 18 | 18 | J1 | 454 | 18 |  | 18 | 16 | J | 523 | 17 |  | 19 |
|  |  |  |  |  |  | J2 | 67 | 9 |  | 1 | 1 |  |  | 22 |  |  |
| K | 562 | 22 |  | 16 | 13 | K | 561 | 22 |  | 16 | 14 | K | 564 | 25 |  | 17 |
| L | 484 | 19 | 2 | 8 | 7 | L | 485 | 19 | 2 | 8 | 7 | L | 582 | 18 | 4 | 11 |
| M | 623 | 18 | 3 | 16 | 12 | M | 622 | 18 | 3 | 16 | 12 | M | 620 |  |  | 16 |
| Mitochondria | 11 | 23 | 2 | 3 | - | Mitochondria | 11 | 23 | 2 | 3 | - | Mitochondria | 11 | 23 | 2 | 3 |
| other contigs | 5 |  | 2 |  |  | other contigs | 13 |  | 1 |  |  |  |  |  |  |  |
| Total | 5294 | 230 | 9 | 144 | 114 |  | 5298 | 230 | 8 | 144 | 115 |  | 5293 | 230 | 6 | 150 |

With these data, we first compared the DSY562 PacBio assembled data (DSY562-HGAP3) with the CBS138 published genome (CGD, version_s02-m04-r02). The DSY562-HGAP3 genome was first annotated with tools available in the software Geneious (version 9.1.8, “live-annotation” tool) with a threshold level of 95% between comparisons. DSY562-HGAP3 was also further annotated on the basis of a recently published annotation using CBS138 and RNAseq data (Linde *et al.* 2015). As observed from Table 3, the number of deduced CDS (locus tag prefix B1J91) in DSY562 was 5294, which was near to the 5293 predicted CDS from CBS138. The DSY565 genome was next annotated on the basis of DSY562 annotations with the same Geneious tool and yielded 5296 CDS (locus tag prefix B1J92). Table 3 indicates that the number of CDS in mitochondrial DNA and numbers of tRNA were identical between strains. rDNAs were identified in DSY562 and DSY565 and were located at telomeric ends of Chr L and M-J as in CBS138.

The similarity between the total number of CDS in the three strains (5293-5298 CDS) obscures however the following divergence: when comparing the CDS of each isolate, only a total of 5225 CDS were shared between them. Sixty five (65) CDS from CBS138 were not detected in DSY562 and DSY565 (**File S3**, Fig. 3). Seventy nine (79) CDS from DSY562 and DSY565 were not found in CBS138. From these 79 CDS, 60 were common to DSY562 and DSY565, thus highlighting unique CDS for each of the strains. These above-mentioned results suggest that several selection events resulted in the loss and gain of several genes between these strains. When inspecting the identity of these 60 genes, we found that most the genes gained in DSY562 and DSY565 as compared to CBS138 were belonging to adhesin-like genes (33). Nine genes originating from DSY562 and DSY565 (named here Tkp5-1 to -9) were most similar to the Tkp5 protein from *Vanderwaltozyma polyspora.* This type of protein encodes elements of Ty-like retrotransposons (see Discussion). Other common genes between DSY strains underwent duplication events including B1J91/B1J92_D02794g (endoplasmic reticulum membrane protein), B1J91/B1J92_M05115g (a putative retrotransposon protein), B1J91/B1J92_H04257g and B1J91/B1J92_H04279g (encoding copper metallothionein MT-II and MT-IIB, respectively). Interestingly, B1J91/B1J92_H04257g and B1J91/B1J92_H04279g were found in four tandem repeats on Chr H as opposed to CBS138, in which only 1 repeat is found. The remaining 10 genes out the 60 common to DSY562 and DSY565 were ranked as pseudogenes in the current CGD (some of which are also described as adhesin-like genes), thus explaining their absence from the CBS genome used in our analysis. Genes that were CBS138-specific (65) included among others adhesin-like genes (6), CAGL0B01243g (MATalpha1) and CAGL0G07175g (RNA binding protein with RNA-directed DNA polymerase activity). Noteworthy is that MATalpha1 is still present in DSY562 and DSY565 but as another CDS (B1J91/B1J92_B00242g). Most of the remaining CBS138-specific genes were of unknown function (48) (see details on **File S3**).

**Figure 3.**
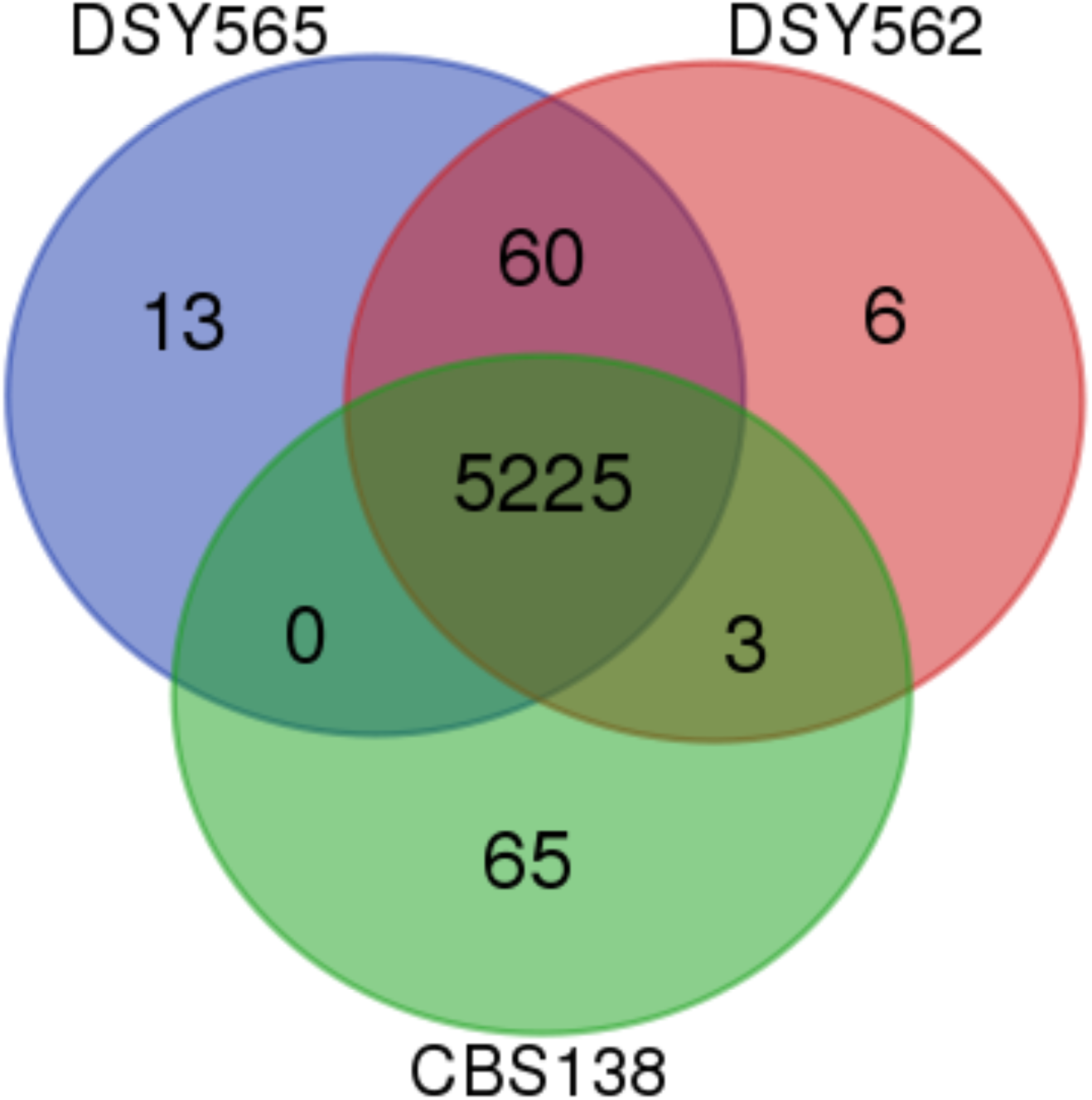
Venn diagram representation of genes shared between DSY562, DSY565 and CBS138. The diagram was generated with a web tool available at http://bioinformatics.psb.ugent.be/webtools/Venn/.

At the level of single nucleotides, large differences were observed between DSY562 and CBS138 (Table 4). Nuclear and mitochondrial genomes diverged by 0.7% between the two strains. These data indicate extensive divergence between the two genomes, however they are in general agreement with the analysis deduced from other *C. glabrata* genomes (Singh-Babak *et al.* 2012). The high number of non-synonymous SNPs (11014) underscores a large number of amino acid substitutions. A high number of SNPs were occurring in genes encoding adhesins, and the following section will discuss the resolution of this important gene category. A few genes underwent frameshifts resulting in protein truncations. For example, B1J91/B1J92_D00154g (*AQY1*, role in ion transport) was truncated in DSY562 and DSY565 (CDS length of 459 bp) as compared to CBS138 (CDS length of 873 bp). Interestingly, another homolog is present in DSY562 and DSY565 (B1J91/B1J92_A01221g), thus likely compensating for the probable loss of function of B1J91/B1J92_D00154g. B1J91/B1J92_C02343g (*ARB1*: protein with ATPase activity and role in cellular response to drug, ribosomal small subunit export from nucleus and cytosol) which contains an intron in CBS138, lacks a reading frame in DSY562 and DSY565. Removal of the intron in this gene restores a truncated CDS as compared to CBS138. B1J91/ B1J92_E01309g (*EKI1*: choline kinase activity, role in phosphatidylcholine biosynthetic process) exhibited a N-terminal deletion due to the use of other intron splicing as compared to the CBS138 homologue. B1J91/B1J92_M05687g (*NPY1*: have NAD+ diphosphatase activity, role in NADH metabolic process), B1J91/B1J92_G03487g (*TMN3*: role in cellular copper ion homeostasis), B1J91/B1J92_I10725g (YGR122W: role in negative regulation of transcription from RNA polymerase II promoter), B1J91/B1J92_E01155g (*RPA14*: RNA polymerase I activity) and B1J91/B1J92_H06017g (*FLR1*: multidrug transporter of the major facilitator superfamily) exhibit N-terminal truncated proteins as compared to CBS138. B1J91/B1J92_H06017g (*FLR1*: multidrug transporter of the major facilitator superfamily). Whether these protein truncations may result in deficient functions remains to be established. We also noticed that some previously annotated tandem CDS in CBS138 were now forming a single CDS in DSY562. For example, B1J91/B1J92_I02816g and B1J91/B1J92_I02838g (*AZF1*: asparagine-rich zinc finger protein) were now generating a single CDS (B1J91/B1J92_I02838g) in DSY562. B1J91/B1J92_E03421g and B1J91/B1J92_E03432g (close homologues of *S. cerevisiae CDA1* and *CDA2* genes, chitin deacetylases) were now forming a single CDS (B1J91/B1J92_E03421g). It is interesting to observe that there are roughly 370000 nt differences between the CBS138 and DSY562 genomes (the latter with the largest genome) thus highlighting that additional/novel genes may be present in the DSY strains (see below). As a last step in these comparisons, we analysed the presence of introns in strains DSY562 and DSY565 as compared to CBS138. This was performed first by transfer of annotations from the recently updated CBS138 genome. We next used RNAseq data from both DSY562 and DSY565 transcriptomes to detect spliced RNAs. Table 4 shows that DSY562 and DSY565 genomes possessed at least 144 intron-containing genes, which is close to the number identified in CBS130 (150). RNAseq data from DSY562 and DSY565 showed that 114 and 115 splice sites, respectively, coincided with the predicted splice sites (**File S4**).

**Table 4:**
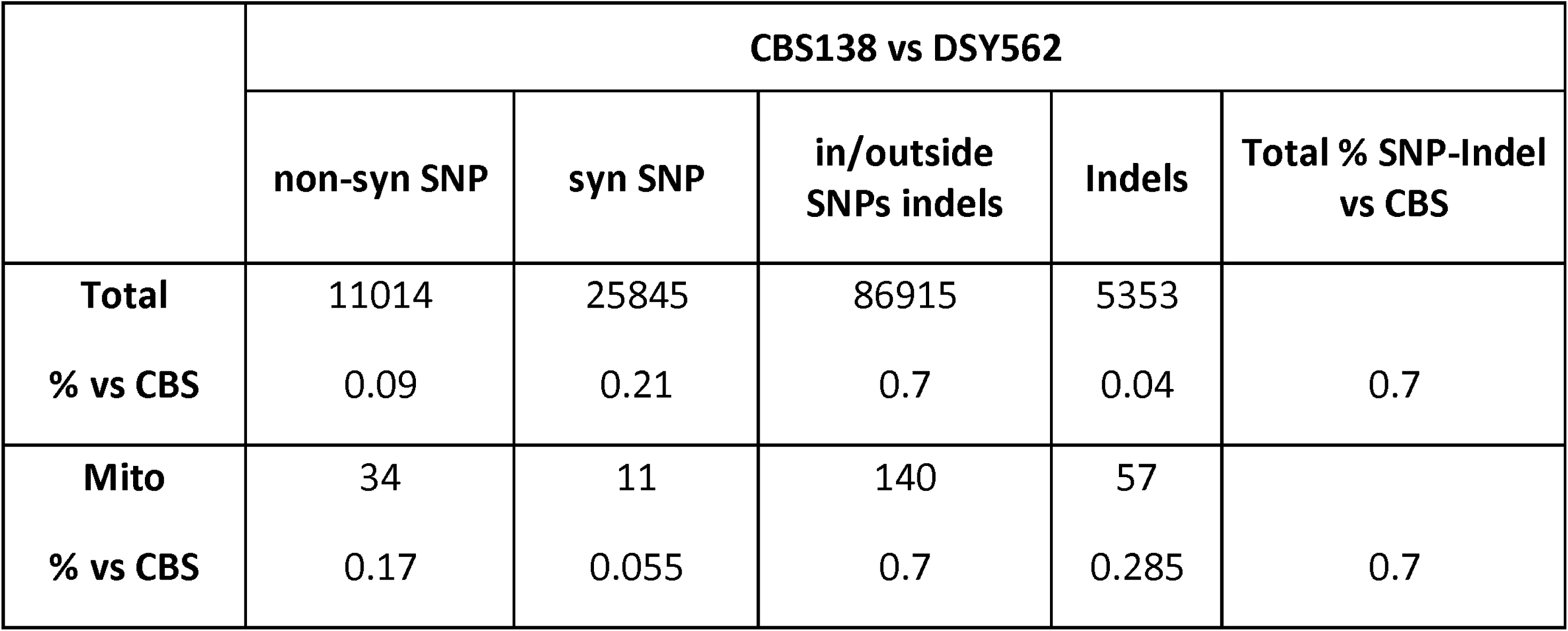
Whole genome differences between DSY562 and CBS138

### The cluster of adhesins in DSY562

*C. glabrata* is known for its extensive repertoire of adhesins. The genes encoding adhesins often contain repeated motifs and are mostly localized at chromosome extremities. These two characteristics make it challenging to establish a detailed and complete listing of adhesins from *C. glabrata* genomes. We have used here PacBio genome assembly approaches to address these challenges and used a systematic analysis (described in Materials and Methods) to identify adhesins-like genes in DSY562 and DS565. Briefly, all DSY562 CDS were screened with the help of the software FungalRV which helps to predict the occurrence of adhesin-like proteins in fungal genomes (Chaudhuri *et al.* 2011). We next analysed the list of potential adhesins for occurrence of PFAM domains and the presence of specific signatures including i) the presence of signal peptides and absence of internal transmembrane domains, ii) the presence of consensus sequence for GPI anchor, iii) the presence of AWP motifs (de Groot *et al.* 2013); iv) the presence of motifs typical for the class of EPA proteins following (Diderrich *et al.* 2015) and (Gabaldón *et al.* 2013), v) the presence of PIR motifs and lastly iv) the occurrence of hyphally-regulated cell wall proteins (Hyphal_reg_CWP, IPR021031). For practical reasons and for comparison purposes with CBS138, we used the same CAGL0 gene naming that is currently accepted for CBS138. When several high similar proteins were identified in DSY562, we append them with a numerical suffix (i.e. 1, 2, etc). According to our analysis, we proposed a list of 101 adhesin-like proteins in DSY562 (**File S5**), which is well exceeding the number of deduced adhesins in CBS138 (66) according to (de Groot *et al.* 2013). Fig. 4 shows the genomic localization of adhesins detected in DSY562 as compared to CBS138. Approximately 50% of the adhesin genes were located in DSY562 near to the telomeric ends of the chromosomes. One example of the expansion of adhesins in DSY562 is given by at Chr C end (Fig 4). The size difference between Chr C between DSY562 and CBS138 may be explained by the tandem arrangement of a group of 4 adhesins. The DSY562 potential adhesins were aligned and a phylogenetic tree was assembled (Fig. 5). This tree was merged with conserved domains features and with CBS138 protein homologues to help on one hand the distinctions between these proteins and on the other hand to better delineate extensions of protein families in DSY562. Several protein clusters (highlighted by colors in Fig. 5) were observed. Members of the EPA family (19) were well discerned in this tree (blue color of Fig. 5) and harbor typical signatures of EPA adhesins including a PA14 domain as well as EPA-specific domains (see details in Material and Methods) with the exception of B1J91_K00170g4. Duplication of specific EPAs occurred in DSY562 for *EPA2, EPA6*, *EPA8*, *EPA12* and *EPA22*. The largest cluster of proteins contained VSHITT motifs repeats and PA14 motifs, however lacked EPA signatures (green color of Fig. 5). CBS138 PWP homologues (PWP1, -2, -3 and -5) were found in this cluster. VSHITT motifs repeats are also found in another cluster (red color of Fig. 5) that essentially lacked PA14 motifs. The largest extension occurred for adhesin-like proteins with VSHITT repeats that include members of the PWP and AWP family (Fig. 5). Another well distinct cluster lacked both VSHITT and PA14 motifs and contained CBS138 AWP homologues (AWP6, -7). Only one gene (B1J91_M11726g) contained a PIR motif and was extensively unrelated to other genes of the phylogenetic tree. The remaining proteins could not be assembled in clusters with common features. Taken together, the data suggest that 61 novel adhesin-like proteins were assigned to DSY562, while 41 remained common with CBS138 (Fig. 5, only 50 adhesin-like proteins without frameshifts were taken into account in these calculations). The majority of the novel proteins exhibited VSHITT domains.

**Figure 4.**
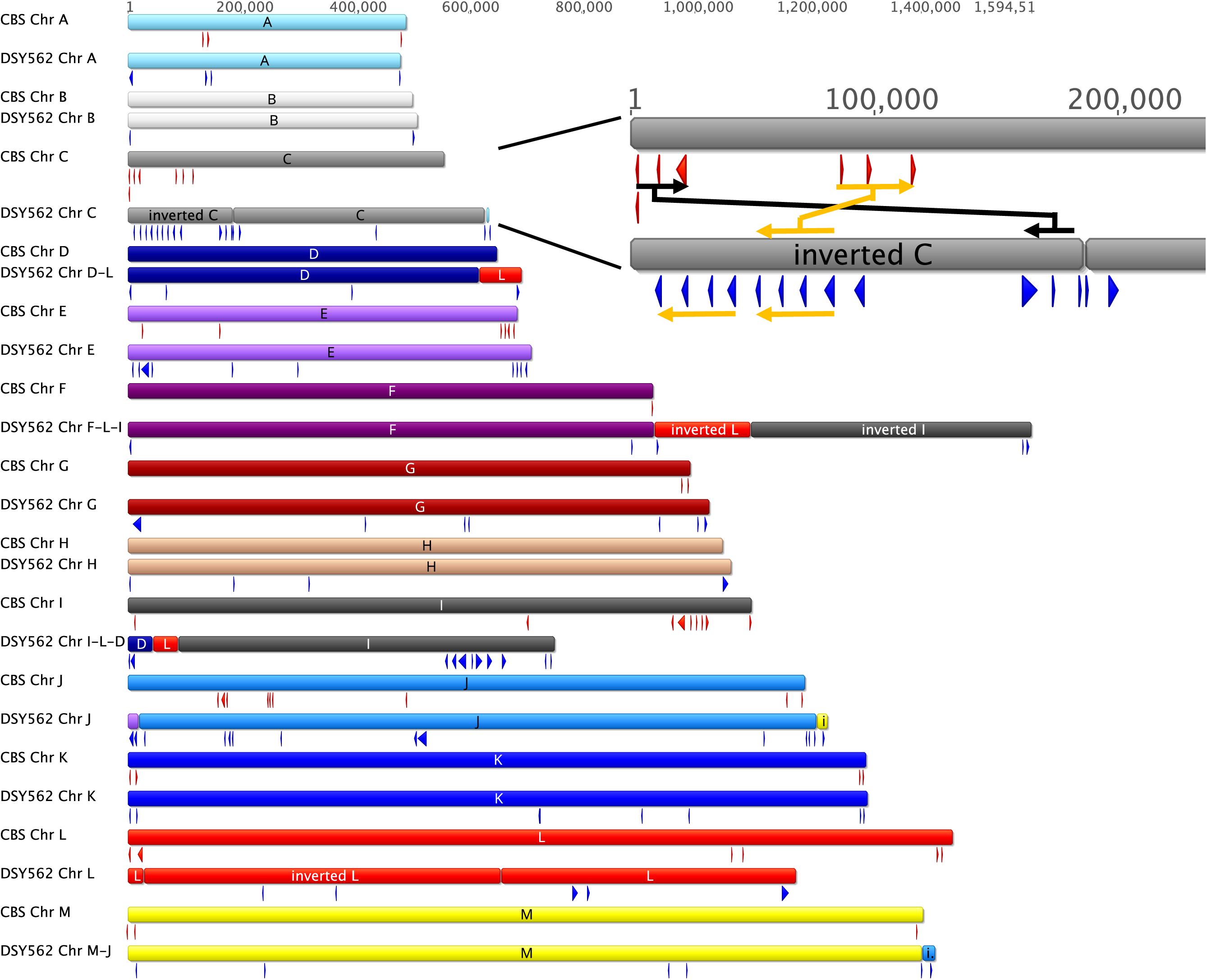
Arrangement of adhesin-like genes in *C. glabrata* CBS138 and DSY562. The position of adhesins is shown by red and blue arrows in CBS138 and DSY562, respectively. Chr C ends of DSY562 and CBS138 were enlarged to highlight the positioning of adhesins in the two genomes. Black arrows indicate the position of adhesins in CBS138 which were inversely located in the Chr C arm of DSY562. A group of 3 tandemly arranged adhesins in CBS138 (orange arrow) is increased by one member in DSY562. This group of 4 adhesins is tandemly arranged with a second group of 4 adhesins.

**Figure 5.**
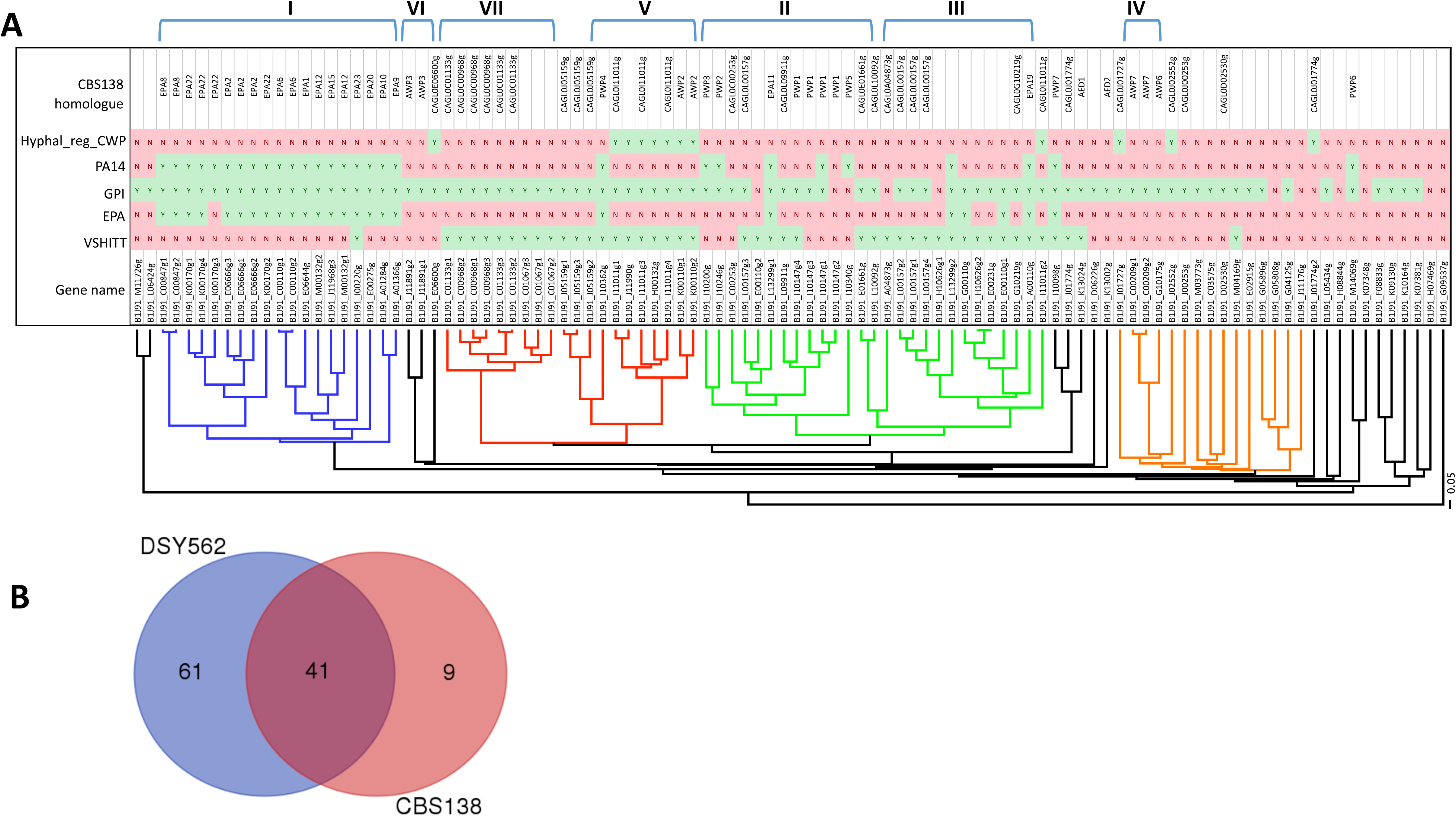
Adhesins in DSY562 Panel A: Phylogenetic tree of DSY562 adhesin-like genes. Each gene is labelled with conserved PFAM functional domains (see Material and Methods). Adhesin clusters (I-VII) were those reported in de Groot *et al*. (de Groot *et al.* 2008). Panel B: Difference between DSY562 and CBS138 adhesins

### Genome comparisons between DSY562 and DSY565

As we mentioned earlier, DSY5652 and DSY565 are two highly related isolates that were obtained from a patient treated with fluconazole for oropharyngeal candidiasis. DSY565 was isolated about 50 days after initiation of fluconazole treatment and was azole-resistant. We reported that azole resistance was due to a mutation in the transcription factor *CgPDR1*. This mutation was participating to an increase of virulence as compared to a parental strain and was measured in different animal models (Ferrari *et al.* 2009; Vale-Silva *et al.* 2016). Given that virulence traits may be multifactorial, we reasoned that other mutations may participate in this phenotype. Direct genome comparisons with PacBio generated data may provide some cues into this hypothesis. We first annotated the DSY562 genome to high resolution as described above and used this basis as a tool to annotate the DSY565 genome. Table 2 and 3 compare several features between DSY562 and DSY565. At the level of the genome, a few changes occurred. While one hundred and one (101) adhesin-like genes were identified in DSY562, 107 adhesin-like genes were found in DSY565. Taking 95 adhesins shared between both isolates (Fig. 6), the discrepancies between the two genomes were due to the presence of adhesin-like genes in small contigs that could not be mapped to the chromosomes. Twelve and 4 adhesin-like and unique genes were identified in the small contigs of DSY565 and DSY562, respectively. In addition, DSY562 possesses 1 gene (B1J91_E00110g1) located at the extremity of in Chr J that is not present in DSY565. This may be due to incomplete assembly of Chr J in this isolate. One other specific change between DSY562 and DSY565 occurred at the left arm of Chr C. While a tandem repeat of B1J91_C01067g is present in DSY562 at the Chr C extremity, only one copy remained in DSY565. However, this gene is now situated between the adhesins B1J92_C00968g1 and B1J91_C01133g1 of DSY565 (Fig. 6). Another major difference between DSY562 and DSY565 is that DSY565 lacks the retrotransposon Tkp5-7 that is situated at the right arm of Chr M in DSY562. Interestingly, DSY565 has retained a single LTR at this locus as a trace of the presence of the retrotransposon.

**Figure 6.**
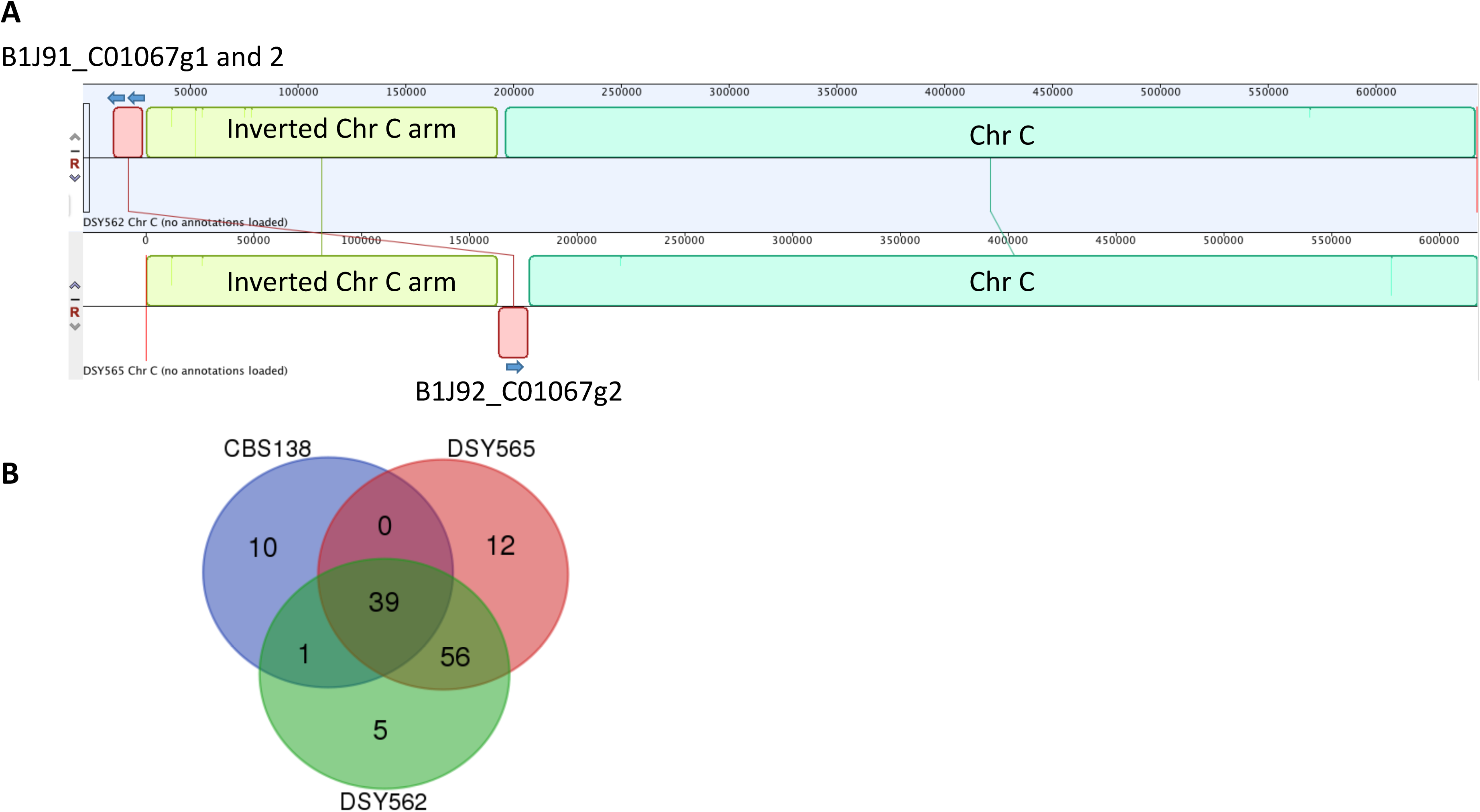
Adhesin distribution between CBS138 and DSY strains. Panel A: Divergent position of adhesins between DSY565 and DSY562. The map was created with the Mauve plug-in available in the Geneious software. Panel B: Venn diagram representation of adhesins shared between CBS138, DSY562 and DSY565 isolates.

At the single nucleotide level, differences between DSY562 and DSY565 include synonymous and non-synonymous SNP variations as well as indels. Table 5 gives an overview of the observed differences. We noticed first that most of the indels in these comparisons were in homopolymeric sequences, which are long stretches (10-30) of identical nucleotides (108 in total). Since it is known that PacBio can yield these differences, we undertook systematic verifications of these sequences by separate Sanger sequencing of small PCR fragments comprising these putative indels. Only 8 of these indels were confirmed by this approach (95 out of 108 were corrected, see Table 5 and File S7). Other indels in non-homopolymeric sequences (14 indels - 8 indels in homopolymeric sequences = 6) were identified by genome comparisons between DSY562 and DSY565. Their presence was confirmed by separate genome alignment with Illumina reads obtained in the early stage of the project (data not shown). We identified also a number of indels within coding regions that were also non-homopolymeric. These variations between the two strains were ranging from 3 to larger number of nucleotides, however only a few of them created frame-shifts (see below in **Table 6B**).

**Table 5:** Nucleotide variations between DSY562 and DSY565

| Chr name | DSY562 vs DSY565 |  |  |  |  |  |
| --- | --- | --- | --- | --- | --- | --- |
|  | non-syn<br>SNP | indels in CDS | syn SNP | in/outside<br>SNP | Indels | corrected<br>Indels in<br>homopolymers |
| **Chr A** | 2 | 0 | 0 | 3 | 0 | 4 |
| **Chr B** | 0 | 0 | 0 | 1 | 0 | 1 |
| **Chr C** | 1 | 1 | 0 | 3 | 1 | 4 |
| **Chr D-L** | 3 | 1 | 1 | 4 | 1 | 7 |
| **Chr E** | 2 | 2 | 2 | 4 | 0 | 8 |
| **Chr F-L-I** | 1 | 1 | 3 | 7 | 3 (2)<sup>a)</sup> | 17 |
| **Chr G** | 0 | 0 | 1 | 1 | 2 (1) | 9 |
| **Chr H** | 1 | 0 | 0 | 2 | 3 | 4 |
| **Chr I-L-D** | 3 | 3 | 1 | 6 | 0 | 4 |
| **Chr J1** | 0 | 1 | 2 | 5 | 3 | 3 |
| **Chr J2** | 0 | 1 | 1 | 0 | 0 | 1 |
| **Chr K** | 1 | 0 | 0 | 1 | 0 | 10 |
| **Chr L** | 0 | 1 | 0 | 9 | 0 | 4 |
| **Chr M-J** | 3 | 0 | 0 | 6 | 1 | 16 |
| **mitochondrial**<br>**DNA** | 0 | 0 | 0 | 0 | 0 | 3 |
| **Total** | 17 | 11 | 11 | 52 | 14 | 95 |

<sup>a)</sup>: number in parentheses indicate the number of single nucleotide indel.

A total of 17 non-synonymous SNPs were identified between both genomes (Table 6A). As expected, the transcription factor *CgPDR1* harbored the L280F substitution in DSY565. This mutation is known to be sufficient to mediate azole resistance in *C. glabrata* (Ferrari *et al.* 2009). The other non-synonymous SNPs were located in genes not known to be associated with drug resistance. Whether or not these mutations are resulting from specific selection pressure from the host or from drug exposure is possible but cannot be answered yet. Besides these non-synonymous SNPs, larger deletions an insertions were identified in coding sequences (Table 6B). The majority of these deletions/insertions (7 out 11) were observed in adhesin-like genes. We cannot presently exclude that these changes are the results of PacBio read miss-assemblies. Some of the affected CDS are located near to chromosome ends, which are known to be recalcitrant to correct assembly and include repetitive sequences. It is likely that recombination events in these regions may generate variations which could result in a beneficial phenotype for the fungus.

**Table 6A:**
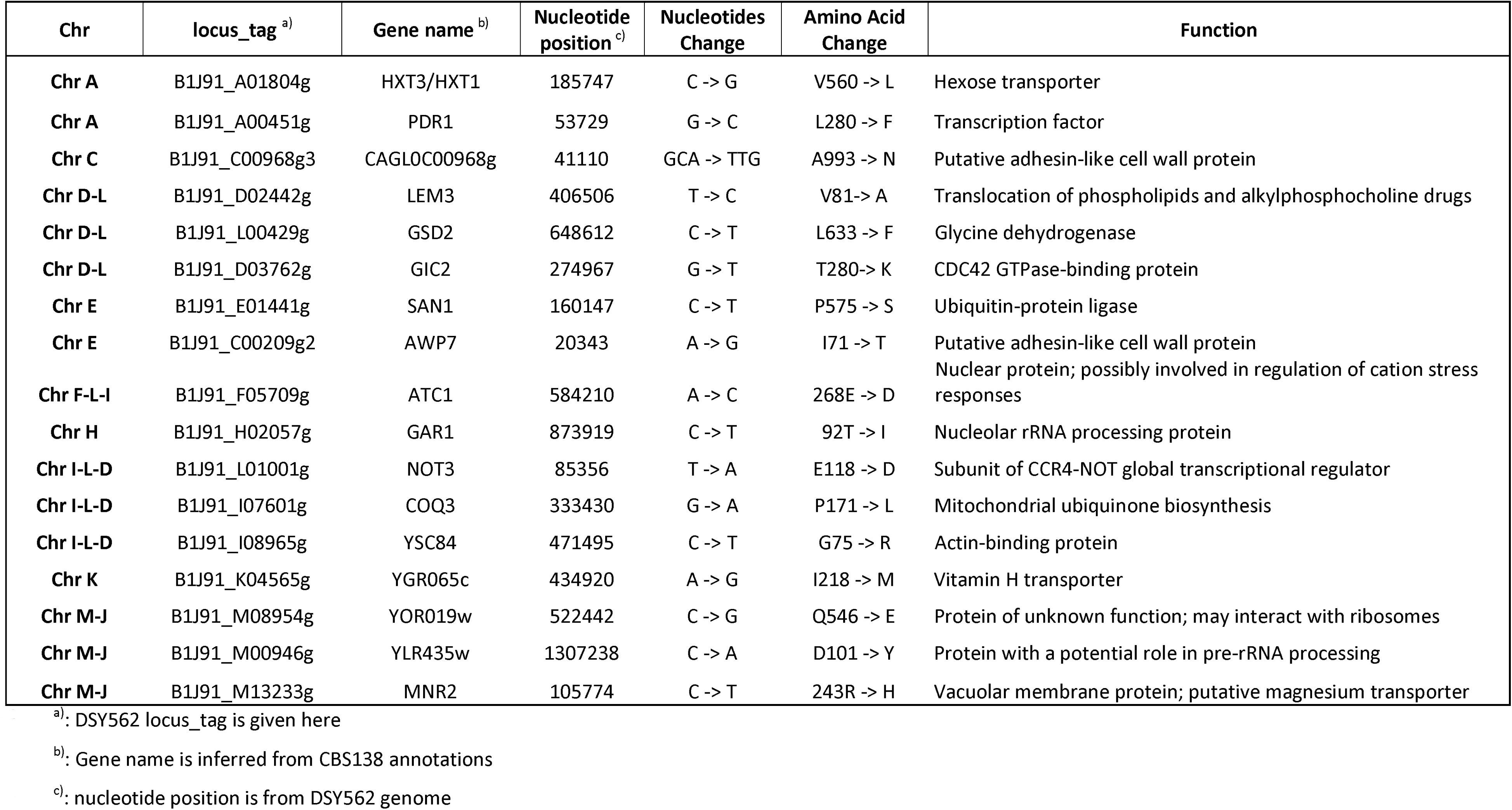
Non-synonymous SNPs between DSY562 and DSY565

**Table 6B:**
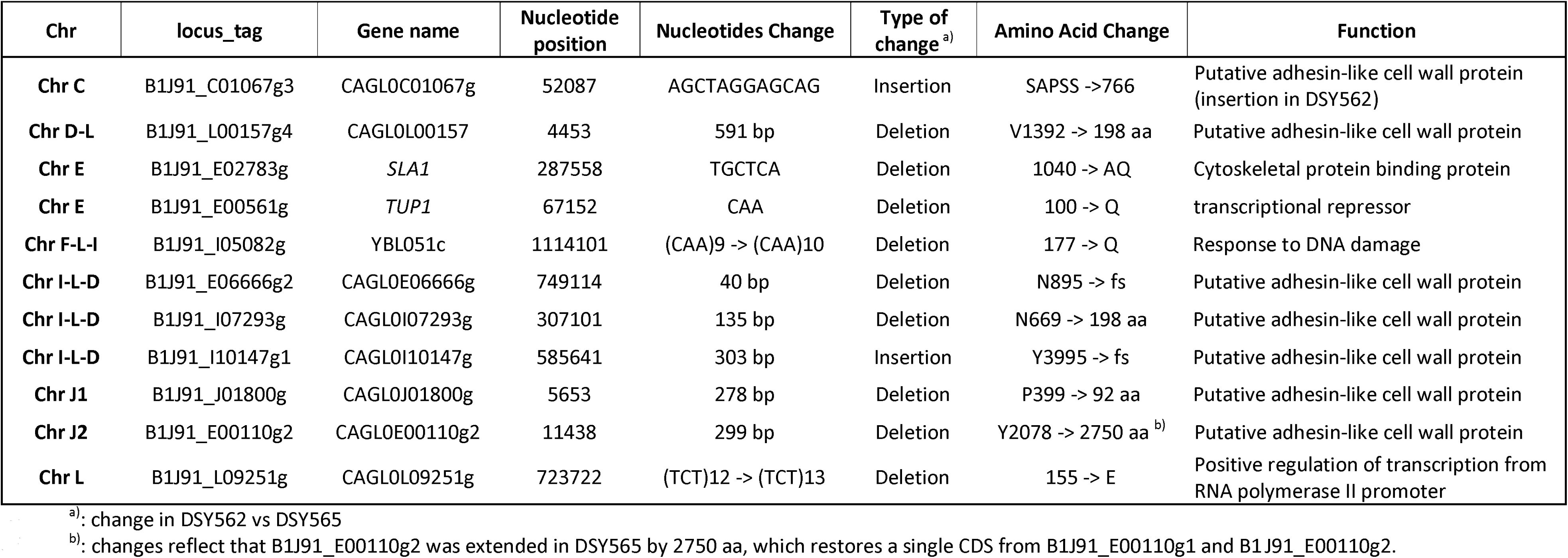
Deletions/insertions in CDS between DSY562 and DSY565

## Discussion

We described here the genomes of two *C. glabrata* sequential isolates, DSY562 and DSY565, originating from a HIV+ patient with oropharyngeal candidiasis. The close relationship between these two isolates reported earlier (Sanglard *et al.* 1999) is confirmed here due to the high level of similarity between the sequenced genomes. We showed earlier that DSY565 was containing a GOF mutation in *PDR1* (L280F) contributing to azole resistance but also to enhanced virulence in animal models. *PDR1* GOF can modulate the expression of the adhesin *EPA1* and this effect results in increased adherence to host tissues. Enhanced virulence due to *PDR1* in DSY565 may also be due to other factors. On the other hand, we observed that DSY562, the parent of DSY565, was intrinsically highly adherent to host cells even with the lowest *EPA1* expression levels as compared to other tested strains. This intrinsically high adherence capacity was observed with a *PDR1* wild type allele (Vale-Silva *et al.* 2016). This suggested the potential presence of other adhesins in this strain background. Given these different reasons, we aimed here to resolve the two *C. albicans* genomes with high resolution in order to identify all possible differences between the two genomes and to determine all possible adhesin and adhesin-like genes in the two genomes.

At the level of the genome, DSY562 and DSY565 underwent several major chromosomal rearrangements. No data are currently available on PacBio-assembled *C. glabrata* genomes of sequential isolates and it is therefore difficult to compare our results with other studies. One recent short report described the genomes of 3 pairs of *C. glabrata* isolates from patients with candidemia, (Håvelsrud and Gaustad 2017). The genomes were assembled de novo using Illumina reads, thus resulting in different genome outputs (119-186 scaffolds). Comparison between the DSY genomes and the three published pairs using same chromosomal fragments highlighted divergences as high as the comparison between DSY isolates and CBS138. This underlines the high genetic diversity existing between *C. glabrata* isolates of different origins which has been observed by others but with different approaches. One other recent work applied also Illumina-based de novo assembly of 33 genomes (Carreté *et al.* 2017), which identified 17 different chromosome patterns. Some patterns are similar to those observed in DSY562 and DSY565. While Chr D-L, Chr L inversions as well as Chr I-L-D are among the common patterns, Chr F-L-I and Chr C inversions seem unique to DSY562 and DSY565. Another separate study established the genome of a *C. glabrata* isolate (CCTCC M202019) which is an industrial yeast strain widely used to produce α-oxocarboxylic acid (Xu *et al.* 2016). Genome data were established on the basis of a 150 bp pair-end library sequenced by Illumina approaches and generated 111 contigs and 74 scaffolds. Interestingly, this *C. glabrata* isolate was remarkably closely related to CBS138 (205 SNPs differences) (Xu *et al.* 2016).

Taken together, these studies underline the high genetic diversity existing between *C. glabrata* isolates of different origins. It is likely that future studies will reveal still additional patterns. In proportion to the size of their genomes, the number of changes between the DSY strains is rather small. We noticed only 17 non-synonymous changes between these isolates (Table 6A). Besides the known L280F change in *PDR1*, several other proteins with diverse functions were affected. It is not immediately obvious to determine the role played by these alterations. One might speculate that they could contribute to the fitness of DSY565 or that they might be accidental. Other larger changes were detected between both strains (Table 6B) and they occurred mainly in adhesin-like genes. This could reflect some adaptation due to host selection pressure.

It has been reported earlier that the mismatch repair gene *MSH2* can modulate drug resistance when its activity is compromised by specific SNPs giving rise to an hypermutator phenotype (Healey, Zhao, *et al.* 2016). Interestingly, DSY562 and DSY565 carry a V239L non-synonymous mutation in *MSH2*, which is known to favor this hypermutator phenotype (Healey, Zhao, *et al.* 2016). One might expect that this phenotype could generate genetic diversity within the context of host conditions. DSY565 was isolated with a time-lapse of 50 days in the oral cavity of a patient as compared to DSY562. Within this elapsed time, the *MSH2* defect may lead to mutation accumulations. In a study probing the accumulation of mutations due to *MSH2* defects in *S. cerevisiae* over 170 generations *in vitro*, mutation rates (mutations per bp per generation) were in the range of 5-9 10^-8^, which corresponded to 8-16 single base pair substitutions and 110-180 deletions/insertions in the haploid *S. cerevisiae* genome (Lang *et al.* 2013). The number of generations that occurred within the time lapse of DSY562 and DSY565 isolation may correspond to at least 250-500 generations, when taking into account 5-10 generations per day according to published data (Forche *et al.* 2009; Roetzer *et al.* 2011). Taking into account data of Table 6A and **B**, the number of SNPs and indels between DSY562 and DSY565 are in agreement with numbers reported in *S. cerevisiae*. One may expect that the *MSH2* defect may generate in *C. glabrata* random mutations with negative effect on *in vivo* fitness. The host conditions may serve as a selection pressure to eliminate progeny with fitness costs. We only sampled successful progenitors from the clinic, which of course strongly reduces the diversity of isolates for further analysis. This underpins that genetic diversity by the *MSH2* defect may be much higher than observed from the collected clinical strains. The current literature on the effect of DNA mismatch repair defects also highlighted that occurrence of mutations is not random: DNA regions with a greater density of repeats are more mutable in mismatch repair defective cells (Ma *et al.* 2012; Lang *et al.* 2013). In the context of adhesins, which are enriched in repetitive elements, *MSH2* defects may perhaps contribute to increase the diversity of this gene family over generations. It has been observed that 44-50% of investigated isolates (n=625) were carrying *MSH2* defects and thus indicate that *MSH2* defect may confer some selective advantage in *C. glabrata* (Healey, Zhao, *et al.* 2016; Healey, Jimenez-Ortigosa, *et al.* 2016; Dellière *et al.* 2016). In agreement with our hypothesis, it is interesting to mention that CBS138 and CCTCC M202019 do not exhibit *MSH2* defects and contains a limited repertoire of adhesins as compared to DSY562/DSY565.

Besides the above-mentioned SNPs, the DSY562 and DSY565 genomes harbored transposon-like genes not reported previously in *C. glabrata*. These Ty-like retrotransposons genes (named here Tkp5-1 to -9) were most similar to the Tkp5 protein from *Vanderwaltozyma polyspora,* a yeast representing the post-whole genome duplication (WGD) lineage most divergent from *Saccharomyces cerevisiae* (Scannell *et al.* 2007). We observed that, while Tpk5-2, -5 to -9 encoded the same proteins, Tpk5-3 and -4 were different (66% identity) but still similar to the Tkp5 protein from *V. polyspora*. Tpk5-1 shared the C-terminal ends of Tpk5-3 and -4. As expected from retrotransposons, Tpk5-2, -5 to -9 were flanked by the same long terminal repeats (LTRs), while Tpk5-3 and -4 were flanked by other distinct LTRs. Curiously, Tpk5-1 was located next to B1J91/B1J92_M05115g on Chr G, which is described as similar to Ty5-6, a *Saccharomyces paradoxus* retrotransposon. Both genes are flanked by the same LTR. Taken together, these features suggest that DSY562 and DSY565 have acquired retrotransposons from different sources and that these strains have undergone several transposition events. Noteworthy is that DSY565 did not retain Tkp5-7, which is otherwise situated at the right arm of Chr M in DSY562. Given that a single LTR remained at this location in DSY565, this suggests an active transposition system. This retrotransposon was not identified in another location in DSY565.

We determined the presence of more than 100 adhesin-like genes in both DSY strains, which was not yet anticipated from other genome-wide studies. This number exceeds by far the numbers published for CBS138 (50-66), depending on the criteria used for selection) and therefore suggests that an expansion of this gene family occurred in our isolates. A genome report on a separate *C. glabrata* isolate closely related to CBS138 identified 49 adhesin-like genes (Xu *et al.* 2016). Since no equivalent *C. glabrata* genome assembly has been yet published using a PacBio approach, we can still not confirm whether or not the investigated isolates constitute a unique case or that a high repertoire of adhesin genes (>100 genes) is shared in several *C. glabrata* clades. To help the distinction of adhesins in DSY562 and DSY565, we used a combination of hierarchical clustering and mapping of PFAM and functional domains. With this approach, we could distinguish 4 major groups including the EPA-like genes, two VSHIT-containing groups and another group with genes lacking EPA- and VSHIT signatures. This grouping is less complex than proposed earlier using CBS138 data (7 clusters) (de Groot *et al.* 2008). The overlay between both grouping showed consistently that cluster I contained the EPA-like genes, that clusters V and VII and clusters II and III were grouped in two categories that include VSHIT-containing adhesins. The other clusters (IV and VI) are located in distinct branches, the latter being grouped in genes lacking EPA-and VSHIT signatures. It is presently difficult to validate one or the other categorization, since no systematic functional analysis of all adhesins has been yet carried out with the exception of EPA-like genes (Zupancic *et al.* 2008). Now that a more extensive repertoire of adhesins is available from our studies, such analysis may be undertaken in the future.

## Acknowledgements

This work was supported to a grant from the Swiss National Research Foundation 31003A_146936. The authors are thankful to F. Ischer for technical assistance and the LGFT (University of Lausanne) for technical support. The purchase of the Pacific Biosciences RSII instrument at the University of Lausanne was financed in part by the “Loterie Romande” through the “Fondation pour la Recherche en Médecine Génétique”.

## Material and Methods

### Strains and media

Strains DSY562 and DSY565 were reported earlier (Sanglard *et al.* 1999) and were grown in YEPD at 30°C under constant agitation.

### DNA isolation

Genomic DNA from DSY5652 and DSY565 was obtained by spheroplasting method as previously described (Calabrese *et al.* 2000).

### Multilocus Sequence Typing (MLST)

MLST was performed according to the study of Dodgson *et al*. (Dodgson *et al.* 2003). Sequencing data of individual loci were analysed using an online web tool (http://cglabrata.mlst.net) to attribute sequence type (ST).

### Genome sequencing

In order to produce high quality genomic DNA from *C. glabrata* isolates, overnight cultures (5 ml) were first grown in YNB minimal medium (0.67% yeast nitrogen base [Dicfo] with 2% glucose) at 30°C under constant shaking to obtain 2 x 10^8^ cells for each strain. The DNA isolation protocol followed instructions of the Gentra Puregene Yeast/Bact kit (Qiagen) with slight modifications. First, yeast cell lysis was performed with Zymolyase 100T (3 μg/μl end concentration) for 30 min at 37°C. In addition, after RNAse A treatment of precipitated nucleic acids, phenol/chloroform extractions were carried out according to recommendations issued by Pacific Bioscience (PacBio SampleNet-shared protocol) for the use of phenol/chloroform/isoamyl alcohol. After final precipitation with NH_4_OAc and several washes with 80% alcohol, the genomic DNA was dissolved carefully in a small volume of elution buffer (10 mM Tris-HCl, pH 8.5). Aliquots of 5 μg of extracted, high-quality, genomic DNA was diluted to 150 μL using elution buffer at 30 μg/μl. Long insert SMRTbell template libraries were prepared (20 kbp insert size) according to PacBio protocols. In total, 9 SMRT-cells per strain were sequenced using P4 and P5 polymerase binding and C2 and C3 sequencing kits with 120 (5 first smrtcells) and 180 min acquisition on PacBio RSII. De novo genome assemblies were produced using PacBio’s SMRT Portal (v2.0.0) and the hierarchical genome assembly process (HGAP version 3.0), with default settings and a seed read cut-off length of 6000 bp. PacBio assemblies are available under Bioproject PRJNA374542.

Illumina-based genome sequencing was performed by FASTERIS SA (Plan-les-Ouates, Switzerland). Briefly, 4 μg of each DSY562 and DSY565 genomic DNA was fragmented by nebulization and the genomic libraries were prepared following the procedure described in the “Genomic DNA Sample Prep Kit” (Illumina, FC-102-1003). To obtain inserts of 300bp, a size-selection was performed on agarose gel before a 10 cycles PCR amplification. The resulting libraries were sequenced in a 2x38bp run on a Genome Analyzer IIx using the Sequencing Kit v4 (Illumina). Paired-end reads from Illumina-based genome sequencing are available under Bioproject PRJNA374542.

The final assembled genomes for DSY562 and DSY565 and their annotations are available under accession numbers MVOE00000000 and MVOF00000000, respectively.

### Genome comparisons

The final PacBio assemblies of DSY562 and DSY565 were analysed using Geneious (9.1.8). Genome comparisons were performed on the basis of each chromosomes homolog using the Mauve aligner plugin (progressive Mauve algorithm) (Darling *et al.* 2004). The alignments were next inspected with a SNP discovery tool integrated in the software.

For comparisons between CBS138 and Illumina-based genome sequencing of DSY562 and DSY565, reads were imported in the CLC genomic workbench software (v.9.5.2) (Qiagen). Variant detection was performed from the mapped reads using both probabilistic and quality-based variant detection tools available in CLC.

### RNAseq

RNA extractions from the *C. glabrata* strains DSY562 and DSY565 were carried out as previously published (Ferrari *et al.* 2011). The strains were grown in to logarithmic phase in 5 ml YEPD medium (1% Bacto peptone [Difco], 0.5% yeast extract [Difco], 2% glucose [Fluka]) at 30°C under constant shaking. RNA libraries for RNAseq were prepared with a TruSeq Stranded Total RNA Library Prep Kit (Illumina). The resulting libraries were processed using the Illumina TruSeq PE cluster kit v3 reagents and sequenced on the Illumina HiSeq 2500 system using TruSeq SBS kit v3 reagents. Sequencing data were processed using Illumina Pipeline software version 1.82 to obtain fastq files. RNAseq data are available under Bioproject PRJNA374542.

### Sanger sequencing

Sequencing reactions were carried with a Big Dye Terminator DNA sequencing kit (Thermo Fisher Scientific, Waltham, MA, USA) and analysed with a 3130XL genetic analyzer (Applied Biosystems). Primers used are listed in Table S1. Forward and reverse tag primer tails (5’-CGACGCCCGCTGATA-3’ and 5’-GTCCGGGAGCCATG-3’) were added to each primer pair (labelled by suffixes “-R” and “-F”). PCR were performed on DSY562 and DSY565 DNA with the forward and reverse primers which was followed by sequencing with the tag primers.

### Adhesins predictions

DSY562 ORFs were analysed with the help of the software FungalRV (http://fungalrv.igib.res.in/index.html), which helps to predict the occurrence of adhesin-like proteins in fungal genomes. To select for potential adhesins, a value of 0.5 was set up as a threshold, as recommended (Chaudhuri *et al.* 2011). The list of potential adhesins was further subjected to several additional tests including a search for PFAM domains (http://pfam.xfam.org/search#tabview=tab1) and the presence of specific signatures relevant for the detection of adhesin-like proteins. These signature included i) the presence of signal peptides and absence of internal transmembrane domains using the online programs SignalP and TMHMM (http://www.cbs.dtu.dk/services/), ii) the presence of consensus sequence for GPI anchor ([NSGDAC] – [GASVIETKDLF] – [GASV] – X(4,19) – [FILMVAGPSTCYWNQ](10)>) as recommended by (Weig *et al.* 2004), iii) the presence of AWP motifs (VSHITT) (de Groot *et al.* 2013); iv) the presence of EPA motifs (EPA CC: CX(27)CX(40)CX(44)DDX(13)CC) and EPA: GCSX(8,9)GL) following (Diderrich *et al.* 2015) and(Gabaldón *et al.* 2013); v) the presence of PIR motifs (Q[IV]XDGQ[IVP]Q) following (de Groot *et al.* 2008).

### Protein alignments

Protein alignments were performed with a webtool available at http://mafft.cbrc.jp/alignment/server (MAFFT version 7) and default settings. A phylogenetic tree was constructed from the produced alignment with the tool The produced alignments were next visualized after average linkage (UPGMA) and a bootstrapping with 1000 resamplings.

### Detection of splicing junctions

RNAseq data obtained from DSY562 and DSY565 were imported as fastq file into Geneious (9.1.8). Splice junctions were detected using Tophat RNAseq aligner (Trapnell *et al.* 2009).

### Counter-clamped homogeneous electric field (CHEF) analysis

Two or three colonies of each yeast strain were inoculated in YEPD liquid medium and incubated overnight at 30°C with agitation. A culture volume corresponding to 10^9^ cells was pelleted at 3000 *g* for 5 min, and cells were washed in 5 ml 50 mM EDTA (pH 9). The pellet was resuspended in 330 μl 50 mM EDTA. Next, 110 μl of SCE (1 M sorbitol, 10 mM EDTA, 100 mM sodium citrate [pH 5.8], adjusted with citric acid) was added and then completed with 5%-mercaptoethanol and Zymolyase 100T (100 units/ml) (solution I). The solution was gently mixed with 560 μl of GTG agarose and rapidly distributed into molds of 10 by 5 by 2 mm. The solidified plugs were immersed in solution II (450 mM EDTA, 10 mM Tris-HCl [pH 8], 7,5% 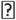-mercaptoethanol) and incubated overnight at 37°C without agitation. Solution II was replaced by solution III (450 mM EDTA, 10 mM Tris-HCl [pH 8], 1% *N*-lauryl-Sarkosyl, and 150 mg/ml proteinase K). Plugs were incubated for 6 h at 65°C without agitation and kept for 10 min on ice. Finally, plugs were transferred in a 0.5 M EDTA (pH 9) solution and stored at 4°C for several months. One-third of the plug was loaded into wells of 0.6% GTG agarose gel in 0.5% Tris-borate-EDTA. The gel was then placed into the electrophoresis chamber of a CHEF DR II (Bio-Rad, Zürich, Switzerland) apparatus. Migration was performed in 0.5% Tris-borate-EDTA at 14°C with the following steps: block 1, 60- to 120-min switch, 6 V/cm, and 120°C over 24 h; block 2, 120- to 300-min switch, 4.5 V/cm, and 120°C over 22 h. After gel electrophoresis, gels were stained with ethidium bromide to reveal DNA under ultraviolet light.

## Supplementary material

**Supplementary File S1:** Comparison of DSY562 with CBS138 based on Illumina reads

**Supplementary File S2:** Comparison of DSY565 with CBS138 based on Illumina reads

**Supplementary File S3**: Characteristics of *C. glabrata* genes common and specific to CBS138, DSY562 and DSY565.

**Supplementary File S4**: Intron predictions and validations in DSY562 and DSY565

**Supplementary File S5**: List of adhesin-like genes in DSY562 and DSY565

**Supplementary File S6:** List of adhesin-like genes for Venn diagram Fig. 6

Supplementary File S7: List of indels corrections in DSY562 and DSY565 genomes

**Table S1:** primers used in this study

